# The Chikungunya virus nsP3 macro domain inhibits activation of the NF-κB pathway

**DOI:** 10.1101/756320

**Authors:** Grace C. Roberts, Nicola J. Stonehouse, Mark Harris

**Affiliations:** School of Molecular and Cellular Biology, Faculty of Biological Sciences, and Astbury Centre for Structural Molecular Biology, University of Leeds, Leeds, LS2 9JT, UK

**Keywords:** Chikungunya virus, nsP3, macro domain, ADP-ribose, NF-κB

## Abstract

The role of the chikungunya virus (CHIKV) non-structural protein 3 (nsP3) in the virus lifecycle is poorly understood. The protein comprises 3 domains. The N-terminus is a macro domain, biochemically characterised to bind both RNA and ADP-ribose, and to possess ADP-ribosyl hydrolase activity – an enzymatic activity that removes ADP-ribose from mono-ADP-ribosylated proteins. As ADP-ribosylation is important in the signalling pathway leading to activation of the transcription factor NF-κB, we sought to determine if the macro domain might perturb NF-κB signalling. We first show that CHIKV infection did not induce NF-κB activation, and could not block exogenous activation of the pathway via TNFα, although TNFα treatment did reduce virus titres. Ectopic expression of nsP3 was able to block TNFα-mediated NF-κB activation and this was dependent on the macro domain, as mutations previously shown to disrupt either ADP-ribose binding or hydrolase activity lost the ability to inhibit NF-κB activation. Lastly, we determined the phenotype of the macro domain mutants in the context of virus infection in a range of cell types. Our data are consistent with cell- and species-dependent roles of the macro domain, however, these phenotypes do not correlate with the ability to inhibit NF-κB activation suggesting that the macro domain plays multiple independent roles in the virus lifecycle.

## Introduction

Chikungunya virus (CHIKV) is an alphavirus of the *Togaviridae* family, and is increasingly becoming a threat to global health. First discovered in Tanzania in 1952, CHIKV is now classified into four distinct genotypes and has spread from Africa to Asia, Europe, and the Americas^1^. CHIKV is transmitted by mosquitos and was previously geographically limited to the distribution of the *Aedes aegypti* mosquito which favours tropical climates. However, mutations in the E1 and E2 glycoproteins, detected in 2004, have enabled spread by the *Aedes albopictus* mosquito, which is more widely disseminated around the globe^2^.

Infection with CHIKV causes Chikungunya fever, characterised by high fever, rash and joint pain which can be debilitating and, in some cases, persistent^3^. CHIKV infection induces a highly inflammatory state, inducing tissue damage^4^. In addition, individuals who have been infected with CHIKV are significantly predisposed to arthritis in later life^5^. In healthy adults, CHIKV is rarely fatal though the risk of death is high in infants, the elderly and the immunocompromised^6^.

The CHIKV genome is a positive-sense, single stranded (ss) RNA. It possesses a 5’ cap and a poly-A tail and contains two open reading frames (ORFs)^7^. The first ORF encodes for the four non-structural proteins, nsP1-4, which are directly translated from the genomic RNA^8^. The second ORF encodes for the structural proteins and is translated from a sub-genomic RNA, generated from a negative sense template following genome replication^9^.

The functions of nsP1, 2 and 4 have been well defined. The capping of viral RNA is performed by the enzymatic nsP1^10^. ATPase, NTPase, helicase, and protease activities are provided by nsP2, the latter being responsible for the processing of the non-structural polyprotein^11,12^. The nsP4 is the RNA-dependent-RNA-polymerase (RdRp)^13^.

Despite numerous studies, the role of nsP3 is still unclear. It is required for RNA replication and interacts with a range of viral and cellular proteins, but the specific role of nsP3 in the CHIKV lifecycle has yet to be defined. NsP3 comprises three domains: at the N-terminus is the macro domain, followed by the alphavirus unique domain (AUD) and the hyper-variable domain (HVD) (Supp. Fig. 1a). The macro domain is found in the proteins of all species, including multiple ssRNA viruses^14^. This domain has been shown to interact with, and hydrolyse, ADP-ribose from ADP-ribosylated proteins. Although it has been shown that macro domains of different species can differ greatly in their affinity for ADP-ribose and enzymatic activity, the CHIKV macro domain has been demonstrated to both bind ADP-ribose^15^ and possess hydrolase activity^16,17^. It has been demonstrated that these two properties of the CHIKV macro domain are essential for virus replication in cell culture^18^. The AUD is a less well understood domain, found to be essential for RNA replication but has also been implicated in virus assembly^19^. The HVD varies greatly in both sequence and size between the alphaviruses and has been more widely studied. This intrinsically disordered and unstructured domain is highly phosphorylated^20^ and facilitates the majority of known protein interactions with nsP3^21^.

Although CHIKV has been shown to induce a robust interferon (IFN) response in infected cells, and this is crucial for survival in infected mice^22^, there has been little characterisation of the other innate immunity pathways in response to CHIKV infection. In particular the interplay between CHIKV infection and the NF-κB pathway remains uncharacterised. A single study demonstrated that CHIKV infection does not activate the NF-κB pathway, but instead was proposed to suppress activation of NF-κB by increasing expression of the micro-RNA miR-146a, which in turn downregulated expression of the upstream modulators TRAF6, IRAK1 and IRAK2^23^.

Interestingly, ADP-ribosylation has been implicated in the signalling of the NFKB pathway, specifically in the formation of the central IKK complex^24^. ADP-ribosylation of NEMO (NF-κB essential modulator, also referred to as IKK-γ) inhibits poly-K63 ubiquitination of the protein, blocking the formation of an active IKK complex and concomitantly preventing activation of the NF-κB pathway. We therefore hypothesised that the CHIKV nsP3 macro domain may be able to perturb NF-κB signalling, as a result of its ability to either bind or hydrolyse ADP-ribose. Consistent with a previous study^23^ we demonstrate that CHIKV infection does not activate the NF-κB pathway, and is unable to suppress exogenous activation of the pathway via TNFα treatment. However, in contrast we further demonstrate that ectopic expression of nsP3 is able to inhibit exogenous activation of the NF-κB pathway, and show that this is mediated via the ADP-ribose binding and hydrolase activities of the N-terminal macro domain.

## Materials and Methods

### Cell culture

Huh7 and A549 cells were maintained at 37°C with 5% CO_2_, in DMEM supplemented with 10% FCS, 0.5 mM non-essential amino acids and penicillin-streptomycin (100 units/mL).

### Virus stocks and infection

Infectious CHIKV used here was derived from the ECSA genotype LR2006-OPY1 isolate of CHIKV and generated and titred as described previously^25^. Cells were washed with PBS, virus diluted to the indicated MOI in serum-free DMEM and applied to cells. Cells were rocked for 5 min, incubated for 1 h at 37°C, prior to removal of virus, washing in PBS multiple times, then media replaced with complete media.

### Plasmid constructs

The expression construct for nsP3 (termed nsP3-F) was generated by PCR amplification of the coding sequence from the DNA virus construct, including flanking restriction sites and a C-terminal FLAG tag sequence, and cloned into pcDNA3.1+ using *BamHI* and *NotI* restriction sites. Macro domain mutants were generated in the nsP3-F construct using site-directed mutagenesis (NEB Q5) (primer sequences available upon request). The GFP construct was generated in a similar manner as the nsP3-F construct but lacking the C-terminal FLAG tag. The FLAG-tagged B14 expression construct was kindly provided by Geoffrey Smith (University of Cambridge) and has been previously described^26^. Constructs were transfected into cells using Lipofectamine 2000 (Life Technologies).

### Luciferase Assays

Cells were transfected with 180 ng of reporter plasmid expressing a Firefly luciferase under the control of an NF-κB sensitive promoter as previously described^27^, alongside 20 ng of transfection control pRL-TK (Promega) using Lipofectamine 2000 (Life Technologies). Luciferase samples were collected and quantified using the Dual Luciferase reporter assay system (Promega).

### Immunofluorescence

Cells were grown on coverslips prior to experimentation. Cells were washed with PBS, fixed with 4% paraformaldehyde for 10 min at room temperature, washed with PBS and permeabilised in 0.5% Triton-X-100 for 10 min, washed with PBS, blocked in 2% BSA for 1 h, washed with PBS and incubated with primary antibodies (rabbit anti-nsP3 kindly provided by Andres Merits, University of Tartu, or mouse anti-p65 Santa Cruz SC-8008) for 1 h. After washing in PBS, secondary antibodies (Life Technologies) were applied for 1 h. Coverslips were finally washed with PBS, rinsed in dH_2_O then mounted in Prolong-gold with DAPI. Cells were imaged using a Zeiss LSM 700 confocal microscope, images were processed using Zen black software.

### Western blotting

Protein concentration of lysed cell samples was calculated using a Bradford assay and 30 μg of protein in Laemmli buffer loaded per well on a 12%, 10%, or 7.5% SDS-PAGE gels for B14, nsP3 and phospho-p105 respectively. Protein were transferred from gels onto membrane (Immobilon FL, Merck) via semi-dry blotter for 1 h at 15 V. Membranes were blocked using blocking buffer (LICOR) for 20 min at room temperature prior to incubation with primary antibody prepared in TBS (rabbit anti-nsP3, anti-phospho-p105 NEB 4806, anti-Flag Sigma F3165, or anti-actin Sigma A1978) rocking overnight at 4 C. Membranes were washed in TBS then secondary antibodies (LICOR) added for 1h at room temperature. Membranes were then washed in TBS + 0.1% Tween, then dH_2_O prior to drying and imaging using the LICOR Odyssey scanner.

## Results

### CHIKV infection does not activate the NF-κB pathway

The NF-κB pathway is activated as part of the innate response to viral infection, leading to expression of antiviral genes, and is frequently targeted for viral immune evasion^28,29^. Given the importance of this pathway it is surprising that the interplay between CHIKV infection and the NF-κB pathway is poorly understood. One study proposed that CHIKV infection led to upregulation of miR-146a, a regulator of expression of NF-κB pathway components, thereby blocking the NF-κB response^23^. As no other studies have been published to date on CHIKV and the NF-κB pathway, we sought to address this gap in our understanding.

To test this, we examined the effect of CHIKV infection on the NF-κB pathway using a reporter plasmid containing firefly luciferase (Fluc) under the control of an NF-κB-sensitive promoter with a second plasmid as a transfection control, renilla luciferase (Rluc) under the control of the constitutive thymidine kinase (TK) promoter. We chose to use Huh7 (human hepatoma) cells for this analysis as they are efficiently infected with CHIKV^30^ and respond well to exogenous (e.g. TNFα) activation of the NF-κB pathway (Fig 1a). Following transfection cells were incubated for 16 h, then either infected with CHIKV (MOI=5) or treated with TNF-α. NF-κB activation was determined by normalising the Fluc values to Rluc. As shown in Fig. 1a, CHIKV infected cells showed no increase in NF-κB activation up to 24 h post infection, indistinguishable from the control (neither infected nor TNF-α treated). In contrast, cells treated with TNFα demonstrated a constant increase in NF-κB activation over 24 h.

**Figure 1.**
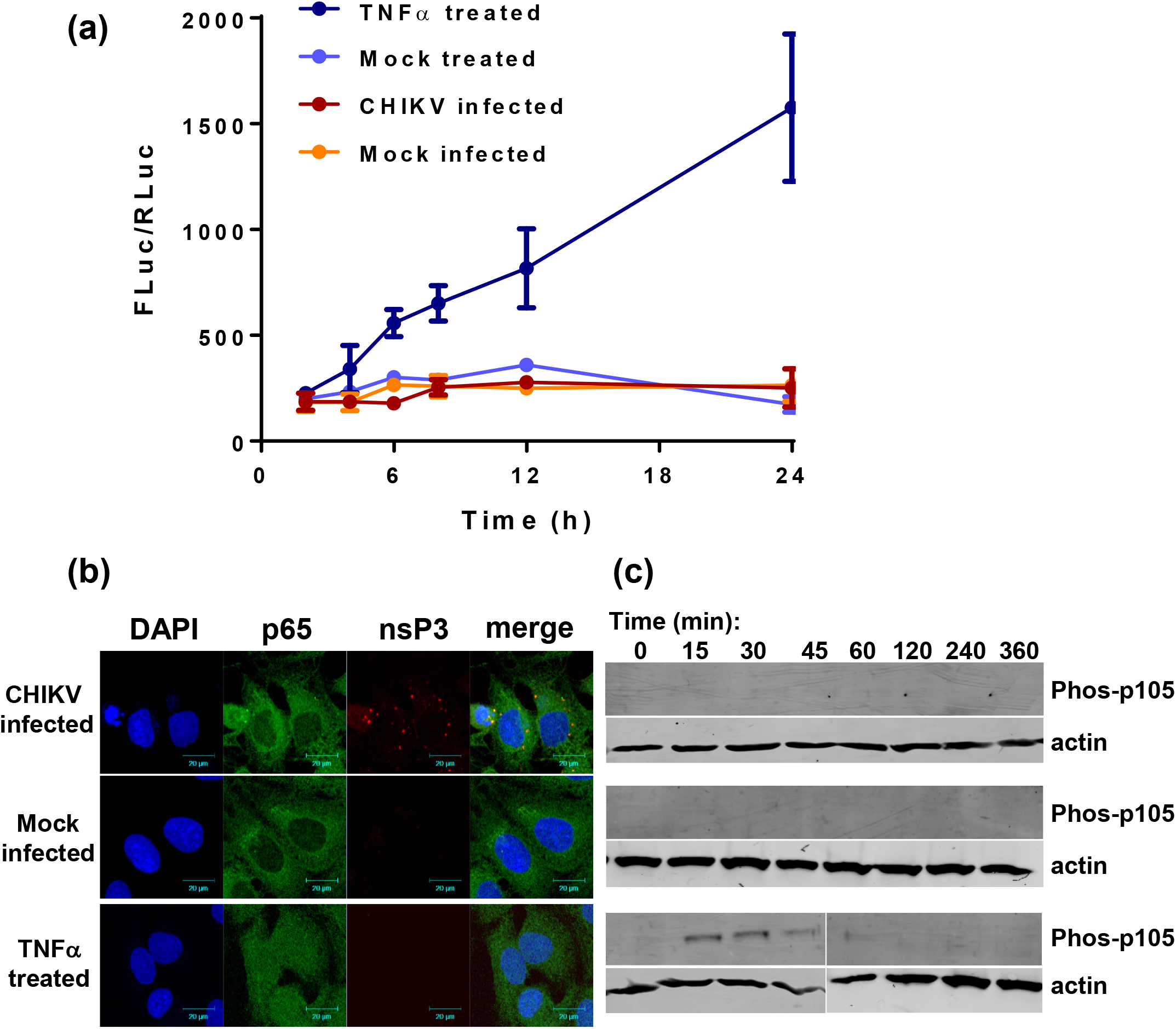
CHIKV infection does not activate the NF-κB pathway. **(a)** Huh7 cells were transfected with NF-κB reporter plasmids 16 h prior to CHIKV infection (MOI=5), mock infection, TNFα treatment (50 ng/mL) or mock treatment. Cells were lysed at indicated time points and assayed for luciferase activity. Firefly luciferase values (Fluc) were normalised to renilla luciferase (Rluc) (n=3). **(b)** Immunofluorescence analysis of NF-κB p65 in infected cells. Huh7 cells were infected with CHIKV (MOI=5), mock infected or TNFα treated (50 ng/mL) fixed at 24 h p.i. and stained for p65 (green) and nsP3 (red). Nuclear p65 indicates activation of the NF-κB pathway. **(c)** Cells were either infected, TNFα or mock treated then lysed at the indicated timepoints. Lysates were western blotted for phospho-p105 with actin as control.

NF-κB activation involves nuclear translocation of the p50-p65 heterodimer. Huh7 cells either treated with TNFα or infected with CHIKV for 24 h were analysed by immunofluorescence using antibodies to p65 to assess NF-κB activation, and nsP3 to detect CHIKV infection. Fig. 1b shows that nsP3-positive (CHIKV infected) cells exhibited a cytoplasmic localisation of p65, consistent with the lack of NF-κB activation following CHIKV infection. As expected, TNFα treatment resulted in translocation of a proportion of p65 into the nucleus of cells, with some protein retained in the cytoplasm.

We then sought to determine whether the pathway was active at an earlier stage. We therefore assessed whether an active IKK complex was able to form in CHIKV infected cells, indicated by the phosphorylation of the NF-κB precursor/inhibitor p105. Western blotting with an antibody to phosphorylated p105 revealed that this was induced by TNFα treatment from 15 min to 1 h.p.t. (Fig. 1c). In contrast, CHIKV infection did not induce the phosphorylation of p105 at any time. These data collectively demonstrate that CHIKV infection does not result in activation of the NF-κB pathway. We proceeded to ask whether exogenous activation of the pathway would have an effect on CHIKV virus production. Treatment of infected Huh7 cells with TNFα resulted in a significant reduction (10-fold) in virus production, as determined by plaque assay titration of virus released into the culture supernatant (Fig. 2a). The data shown in Fig. 2b confirm that TNFα treatment of CHIKV infected cells resulted in p65 nuclear translocation of NF-κB, consistent with NF-κB activation. These data demonstrate that exogenous activation of the NF-κB pathway can inhibit the CHIKV lifecycle and reduce production of progeny virus.

**Figure 2.**
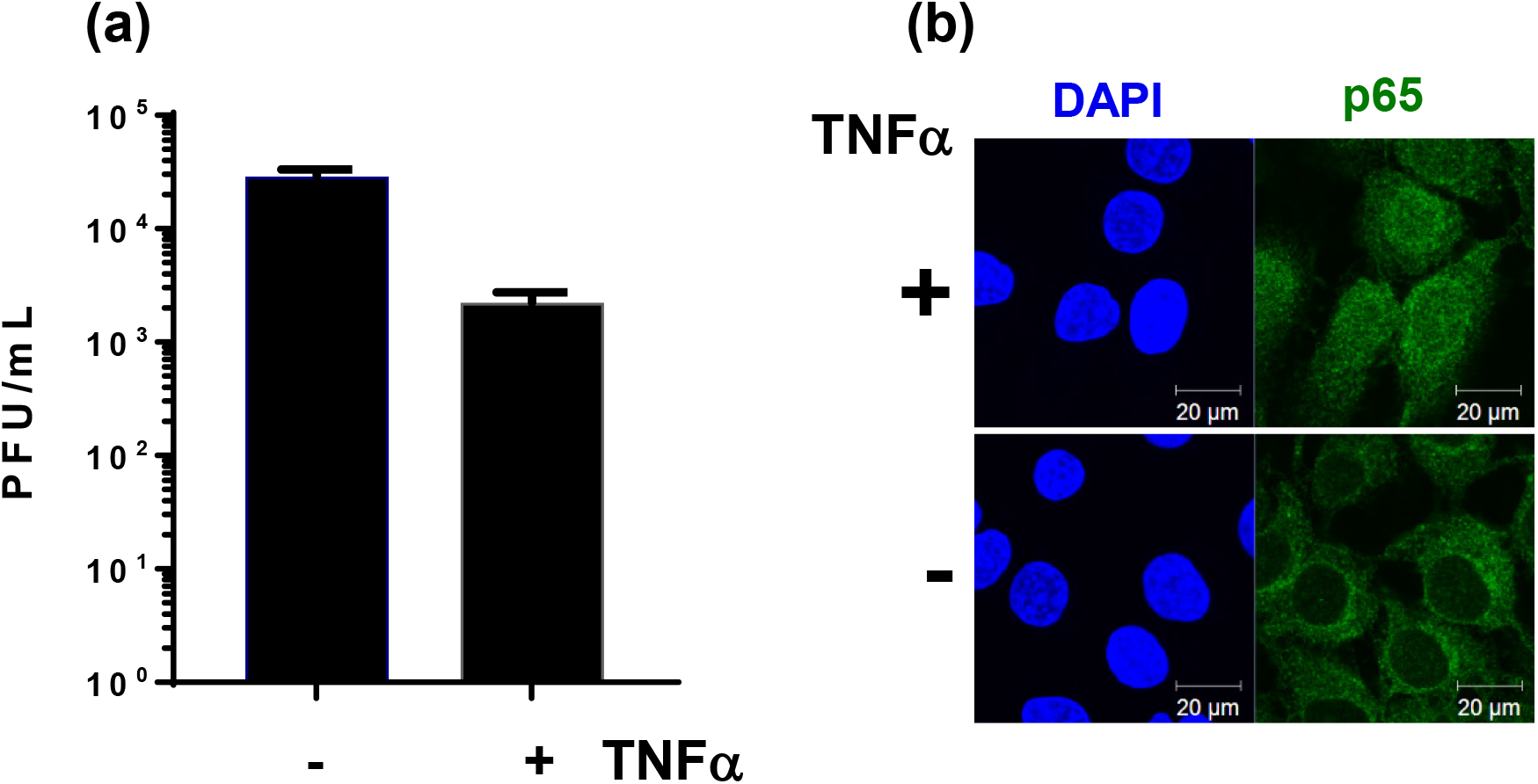
Activating the NF-κB pathway reduces CHIKV titres. **(a)** Huh7 cells were infected with CHIKV at MOI=5, in the absence or presence of TNFα (50 ng/mL). Supernatant was harvested at 24 h for each sample and titred by plaque assay (two separate experiments combined, each at n=2). **(b)** In parallel to (a), cells were TNFα or mock treated, fixed at 6 h p.t. and immunostained for p65 to confirm activation of the NF-κB pathway by TNFα treatment.

### CHIKV infection does not inhibit exogenous activation of the NF-κB pathway

Given that NF-κB activation inhibited CHIKV virus production, we sought to investigate whether CHIKV infection had any effect on exogenous activation of the NF-κB pathway. To test this, Huh7 cells were transfected with the NF-κB reporter plasmid and infected with CHIKV at 16 h post-transfection. Immediately after infection, cells were treated with TNFα, and cell lysates were harvested over a 24 h time period. This analysis revealed that CHIKV infection was not able to inhibit the exogenous activation of NF-κB (Fig. 3a). To confirm this result, cells were analysed by immunofluorescence for nsP3 and p65. As shown in Fig. 3b, CHIKV infection did not block the nuclear translocation of p65 following TNFα treatment. Taken together, these data demonstrate that CHIKV infection is unable to suppress exogenous activation of the NF-κB pathway by TNFα.

**Figure 3.**
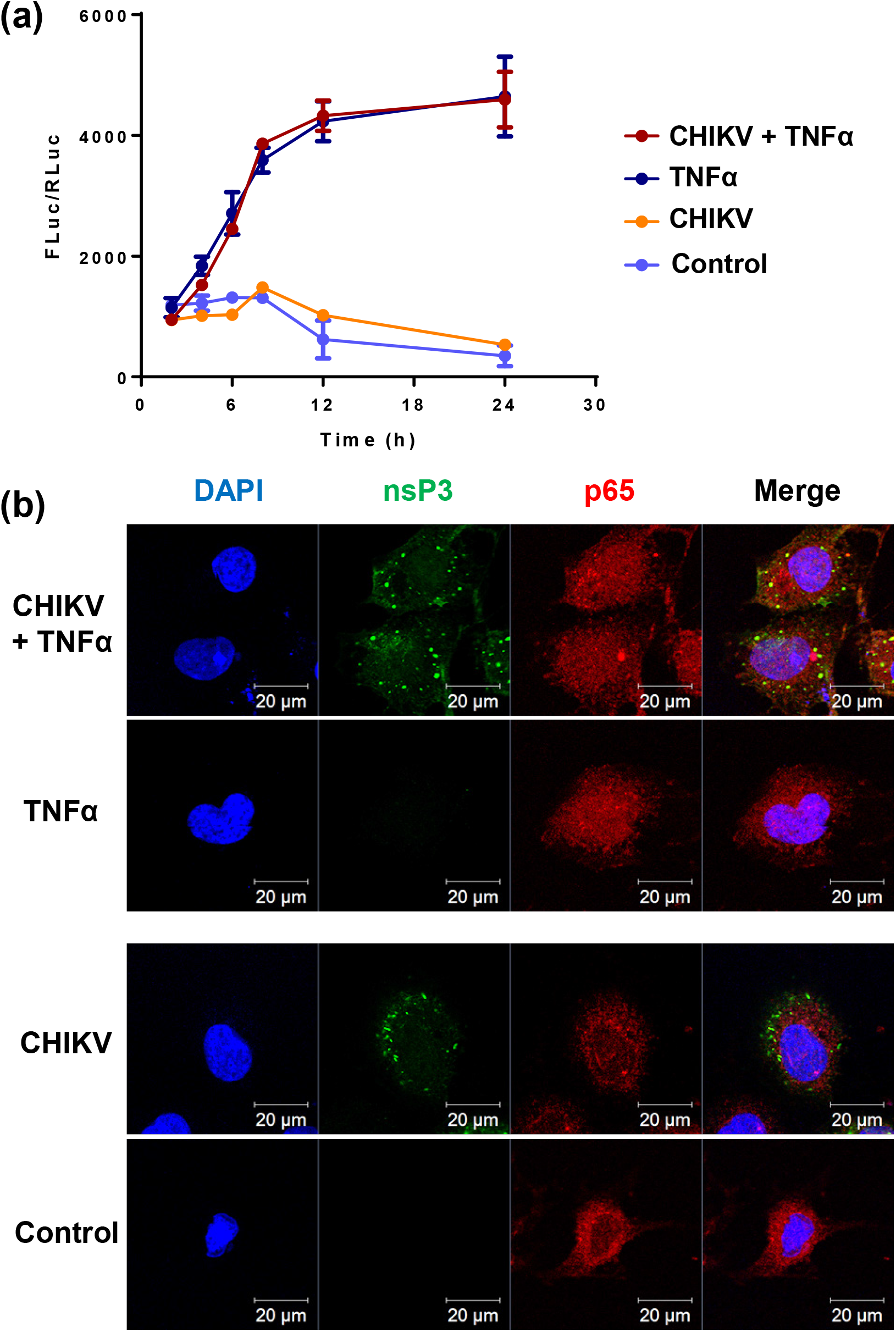
CHIKV infection does not inhibit TNFα-induced NF-κB activation. **(a)** Huh7 cells were transfected with the NF-κB reporter plasmids, incubated for 16 h prior to either infection with CHIKV (MOI=5) or mock infected for 1 h, then either TNFα treated (50 ng/mL) or mock treated. Cells were lysed over a 24 h period and lysates assayed for luciferase. Data shown as Fluc signal normalised to Rluc (n=3). **(b)** Huh7 cells were fixed at 12 h p.i. and immunostained for nsP3 (green) and p65 (red), with a DAPI counterstain for nuclei.

### Ectopic expression of nsP3 in the absence of other CHIKV proteins inhibits the NF-κB pathway

We originally hypothesised that by virtue of its ADP-ribosyl-binding and hydrolase activities, the CHIKV nsP3 macro domain might be able to modulate NF-κB activation. Data presented thus far indicate that in the context of virus infection this was clearly not the case, however it was conceivable that any effect of the macro domain might be masked by antagonistic effects of other viral components. To formally ask whether the macro domain could function to modulate NF-κB activation, we assessed whether any effect could be observed when nsP3 was expressed in the absence of other viral proteins. To do this a C-terminally FLAG-tagged nsP3 expression construct was generated (nsP3-F). As a positive control we utilised an N-terminally FLAG-tagged vaccinia B14 construct (F-B14). B14 is a potent inhibitor of the NF-κB pathway, it binds IKKβ and blocks formation of an active IKK complex^31^. These experiments were conducted in A549 cells as they effectively supported plasmid driven expression of nsP3. Cells were transfected with NF-κB reporter plasmids, together with either nsP3-F or F-B14 expression constructs, with GFP or empty vector as negative controls, incubated for 16 h prior to treatment with TNFα for a further 6 h. As shown in Fig. 4a, both nsP3 and B14 exhibited a similar inhibitory effect on NF-κB activity, both basal levels and the levels of NF-κB exogenously activated via TNFα. The expression of GFP exerted no significant effect on NF-κB activation. Expression of nsP3, B14 and GFP was confirmed by western blot for both nsP3-F and F-B14 (Fig. 4b), and by IF for GFP (Fig. 4c).

**Figure 4.**
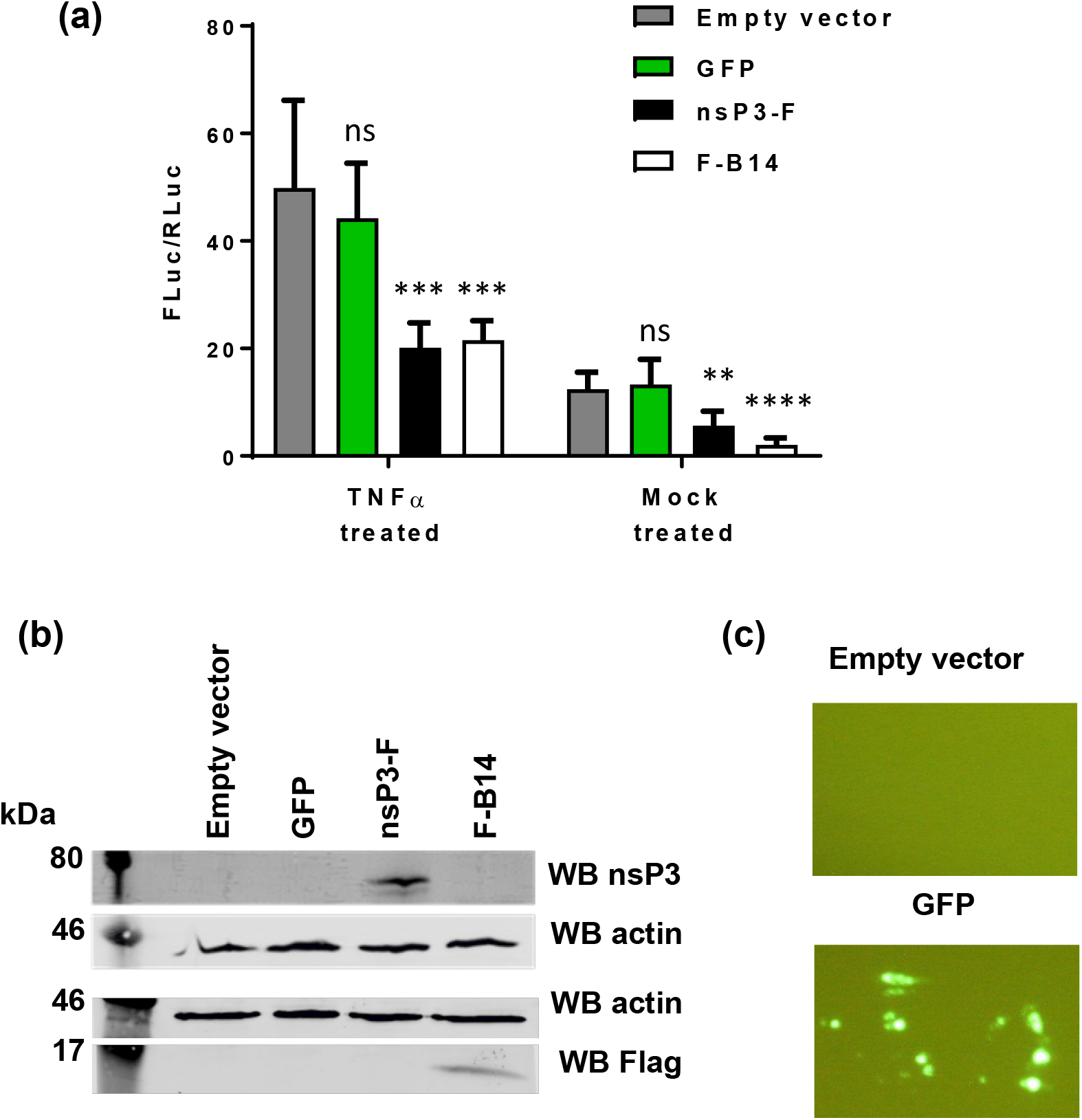
Expression of nsP3 alone can inhibit NF-κB activation. **(a)** A549 cells were transfected with the NF-κB reporter plasmids, and co-transfected with expression constructs for either GFP, FLAG-tagged nsP3 (nsP3-F), FLAG-tagged vaccinia B14 (F-B14) or empty vector. At 16 h p.t. cells were TNFα or mock treated for 6 h prior to lysis, and assayed for luciferase activity (two separate experiments combined, each n=3, data analysed by One-way ANOVA with Bonferroni correction compared to empty vector). ns (*P*>0.05), ** (*P*<0.01), *** (*P*<0.001) or **** (*P*<0.0001). **(b)** Western blot for nsP3-F (59 kDa) and F-B14 (15 kDa). **(c)** Wide field fluorescent microscopy images of GFP-expressing cells compared to cells transfected with empty vector.

To confirm these findings, we again performed immunofluorescence to determine the subcellular localisation of the p65 subunit of NF-κB. As demonstrated in Fig. 5, TNFα treatment resulted in nuclear translocation of p65 in cells transfected with empty vector or GFP. Conversely, p65 was excluded from the nucleus in cells expressing nsP3-F, indicating that nsP3 was inhibiting nuclear translocation of NF-κB, and therefore activation of the pathway.

**Figure 5.**
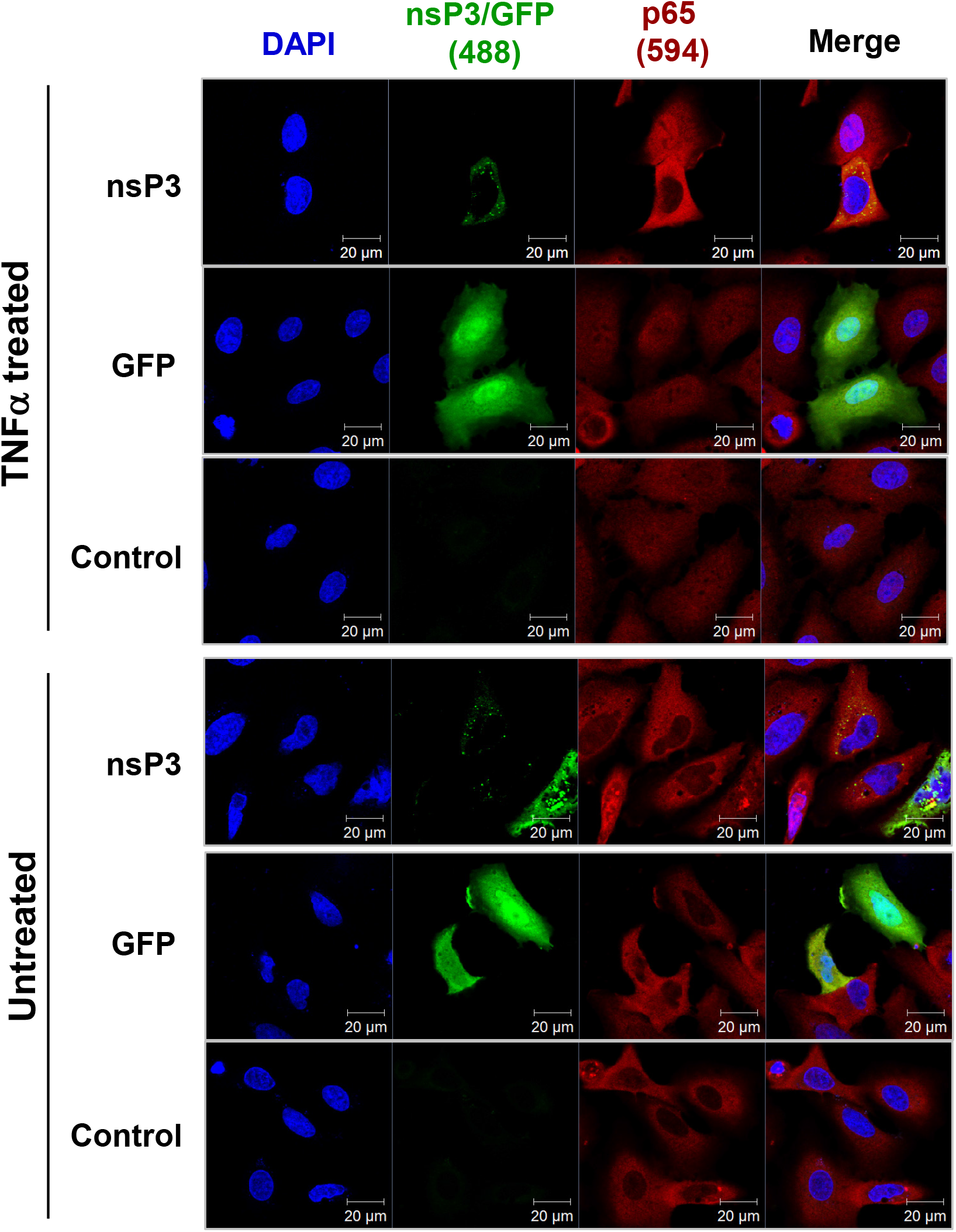
Expression of nsP3 alone blocks nuclear translocation of NF-κB. A549 cells were transfected with either nsP3-F, GFP or empty vector (control) plasmids. At 16 h p.t., cells were treated with TNFα or mock treated for 6 h prior to fixation and staining for p65 (red). Cells were co-stained for nsP3 (shown in green/488), except GFP expressing cells.

### The nsP3 macro domain contributes to inhibition of NF-κB activation

To assess whether the macro domain was involved with the nsP3-mediated inhibition of the NF-κB pathway, we generated a panel of six alanine-substitutions of residues previously shown to be critical for the various functions of this domain. The previously determined phenotypes of these mutants are listed in Table 1 and their locations on the CHIKV macro domain structure illustrated in Supp. Fig. 1b. The panel of mutant nsP3-F expression constructs were transfected with NF-κB reporter plasmids into A549 cells as described above. None of the macro domain mutants could inhibit TNFα activation of NF-κB to the same level as the wildtype (WT), although two mutants (V113A and Y114A) did exhibit a significant inhibitory effect (Fig. 6a). G32A exhibited an intermediate effect, whereas levels of NF-κB activation in the presence of the three mutants D10A, T111A, and G112A were indistinguishable from the negative controls, implicating these residues in the inhibition of the NF-κB pathway by the macro domain. Expression levels for the mutant panel and the B14 control were confirmed by western blot (Fig 6b).

**Table 1:**
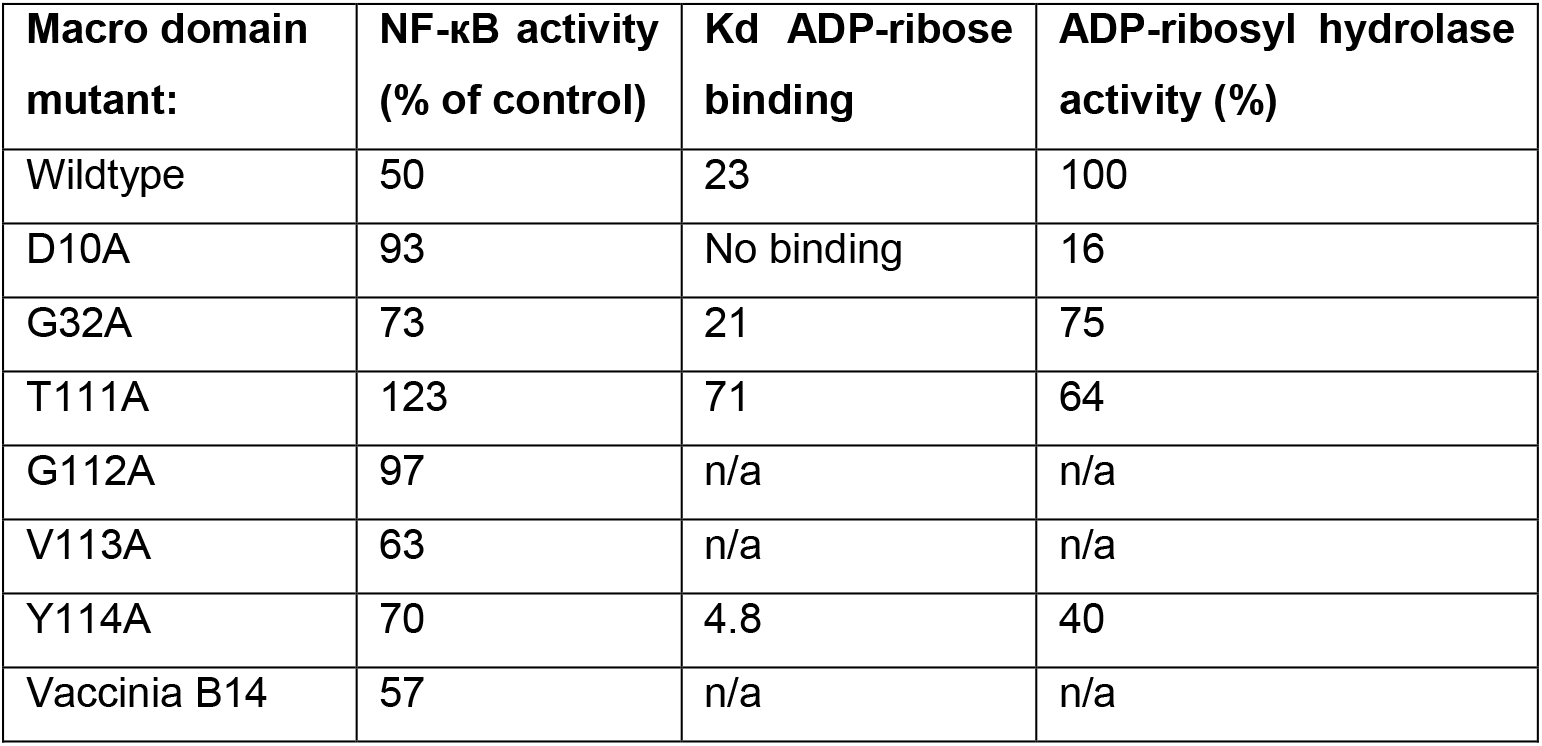
Macro domain mutant phenotypes. Values for ADP-ribose binding and ADP-ribosyl hydrolase activity taken from^18^.

**Figure 6.**
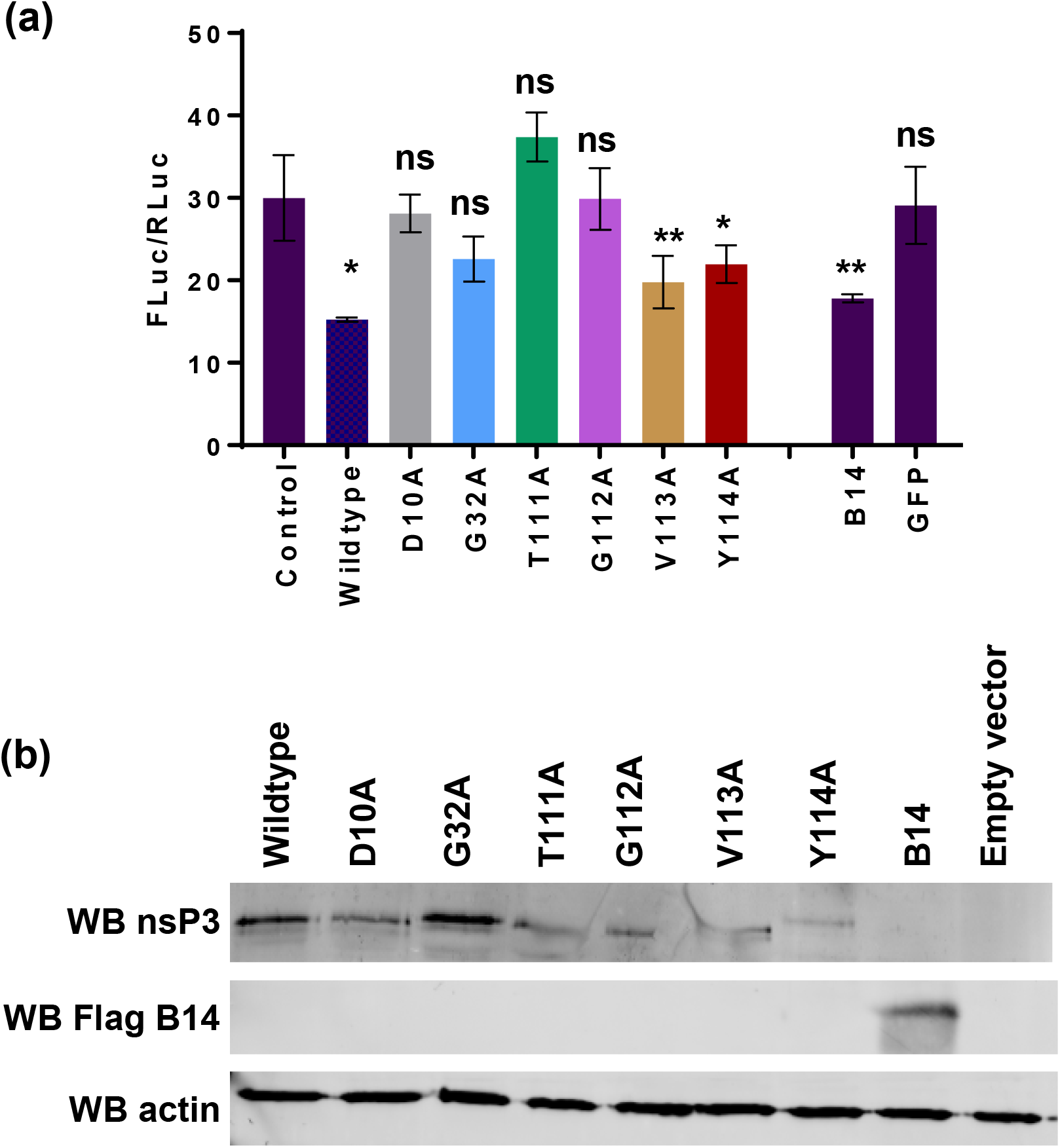
The nsP3 macro domain contributes to inhibition of NF-κB activation. **(a)** A549 cells were transfected with the NF-κB reporter plasmids and either an empty vector (control), WT nsp3-F, the indicated panel of alanine-substitution mutants, F-B14 or GFP. At 16 h p.t., cells were TNFα activated prior to lysis at 6 h post treatment and assay for luciferase (n=3, data analysed by One-way ANOVA with Bonferroni correction compared to empty vector control). ns (*P*>0.05), * (*P* ≤ 0.05), ** (*P*<0.01). **(b)** Confirmation of protein expression by western blotting for nsP3 and Flag tag (for F-B14) on the corresponding cell lysates.

To attempt to correlate the ability of the macro domain to inhibit NF-κB activation with its role in the virus lifecycle we evaluated the phenotype of the mutant panel in the context of infectious CHIKV. We tested the replication of the macro domain mutant panel in 5 cell types: Huh7, BHK, C2C12 (murine myoblasts), U4.4 and C6/36 *(Aedes albopictus).* As a negative control we included an inactivating mutant in the nsP4 RNA-dependent RNA polymerase (GAA). Interestingly the phenotype of the panel varied between the different cell types. In Huh7 cells only two of the mutants (T111A and Y114A) were able to produce any detectable infectious virus (Fig. 7a) and the titres of these were significantly reduced compared to WT. In BHK-21 cells (Fig. 7b), D10A and G112A failed to produce any infectious virus, V113A was attenuated and the other 3 mutants exhibited a WT titre, and similar levels of nsP3 expression detected by western blot (Supp. Fig. 1c). A similar picture was observed in C2C12 cells (Fig 7c), with the exception that D10A was able to produce significant amounts of infectious virus. In U4.4 mosquito cells (Fig 7d) D10A, G112A and V113A failed to produce infectious virus but the other mutants showed no phenotype. Intriguingly, in C6/36 cells (Fig 7e), which lack a functional RNAi response due to a frameshift mutation in the Dcr2 gene^32^, two of these mutants (D10A and V113A) were able to produce infectious virus, suggestive of a role of the macro-domain in counteracting the insect antiviral system. Lastly, when we analysed the phenotype of the mutant panel in the context of a dual-luciferase sub-genomic replicon (SGR)^30^ in Huh7 cells (Fig 7f) we observed that all the mutants were able to replicate to some extent, although only two (V113A and Y114A) retained WT levels of replication. We conclude from these data that the macro domain is involved in multiple stages in the CHIKV lifecycle and exhibits both cell- and species-specific effects. Furthermore, there appears little obvious correlation between the ability to inhibit TNFα mediated activation of NF-κB, and the requirement for virus replication.

**Figure 7.**
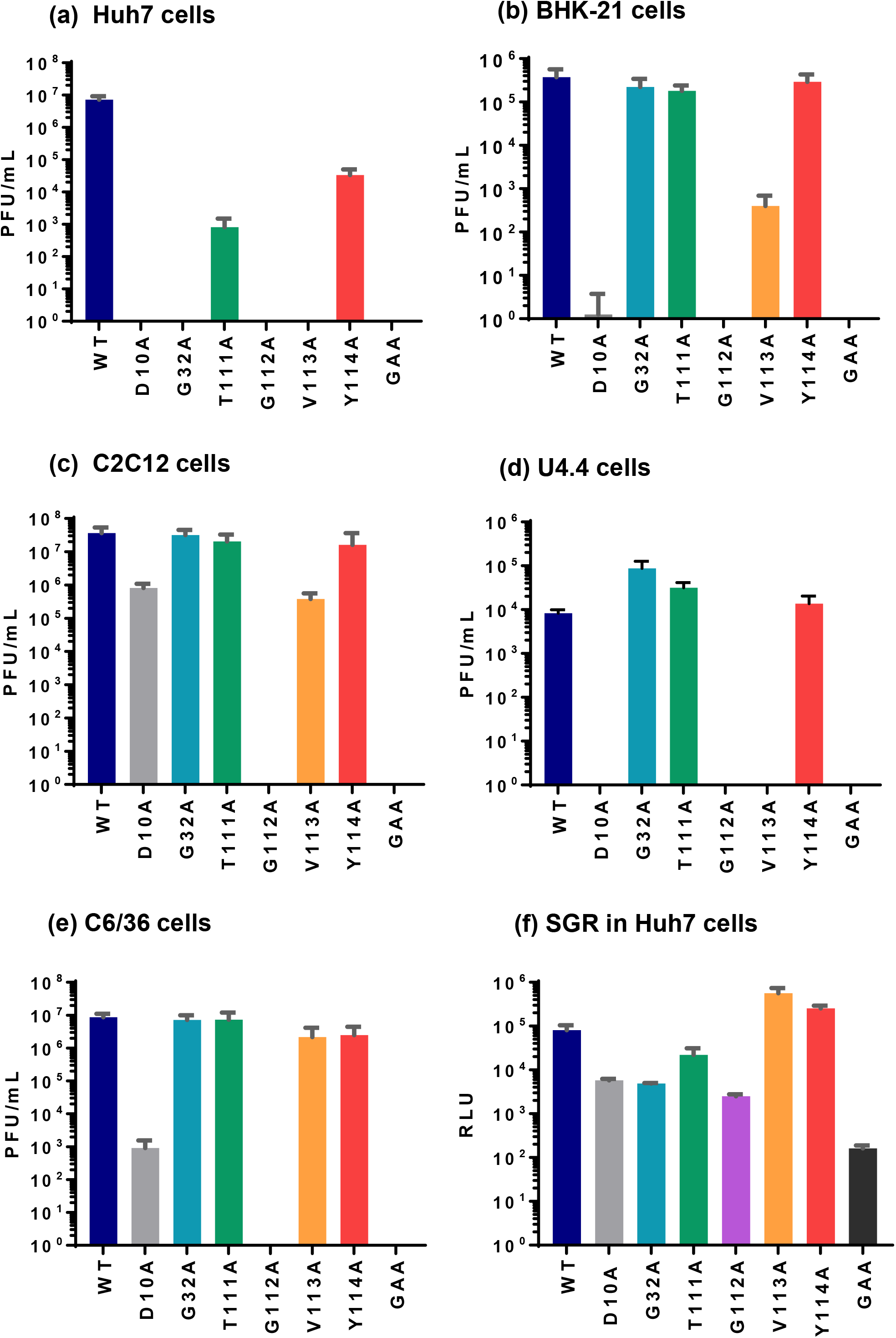
Phenotype of nsP3 macro domain mutants in the context of infectious virus or subgenomic replicon. **A-E** the indicated cell lines were electroporated with either WT ICRES CHIKV RNA or the corresponding panel of macro domain mutants. Virus titres in the supernatant were determined at either 24 or 48 h p.t. by plaque assay on BHK-21 cells. **F.** Huh7 cells were transfected with an ICRES derived dual-luciferase subgenomic replicon RNA (WT or macro domain mutant derivatives) and harvested for luciferase assay at 24 h p.t.

## Discussion

The transcription factor NF-κB is involved in the host cell defence against virus infection and its activity is induced via a variety of canonical and non-canonical pathways^29^, in particular pattern recognition receptors (PRRs) involved in innate immune detection of virus components. Virus infection is a well-characterised inducer of NF-κB activity and most viruses have evolved mechanisms of inhibiting this activation to evade this response, a good example is vaccinia virus which expresses no less than 9 different proteins that antagonise the NF-κB pathway^33^.

In this study we sought to investigate the effect of CHIKV infection on NF-κB activity. We observed that CHIKV infection could neither induce the pathway, nor inhibit activation via the canonical pathway as a response to the external stimulus provided by TNFα treatment. However TNFα treatment did result in a reduction in CHIKV virus production. We reasoned that the lack of NF-κB activation was consistent with the hypothesis that CHIKV was able to block this activation. A likely mediator of this effect was the macro domain of nsP3, and indeed ectopic expression of nsP3 and a panel of mutants in the active site of the macro domain revealed that this was indeed the case. As shown in Table 1 those mutants previously shown to abrogate ADP-ribosyl hydrolase activity (in particular D10A) also abolished the ability to inhibit NF-κB activation, However, this was not an absolute correlation and in fact all mutants had some effect, suggesting that the ability of the macro domain to block NF-κB activation is not solely due to hydrolase activity.

At this stage in our investigation we do not know at which point nsP3 is antagonising the NF-κB pathway, however, as CHIKV infection does not induce the phosphorylation of p105 or the translocation of p65 into the nucleus this provides some clues. Phosphorylation of p105 is mediated by an active IKK complex, thus one attractive hypothesis is that the nsP3 macro domain is perturbing the ADP-ribosylation of NEMO, a key component of the IKK complex. Consistent with this a cellular mono-ADP-ribosyltransferase, ARTD10 (PARP10), inhibits TNFα mediated NF-κB activation by interfering with poly-K63 ubiquitinylation of NEMO^24^. Indeed both NEMO and ARTD10 were shown to be *in vitro* substrates for the ADP-ribosylhydrolase activity of the nsP3 macro domain^17^, and were efficiently de-mono-ADP-ribosylated (deMARylated). Although we have been unable to biochemically demonstrate an interaction between nsP3 and either NEMO or ARTD10, we did observe partial colocalisation of nsP3 with ARTD10 at late stages of CHIKV infection (Supp. Fig. S2). This is consistent with the hypothesis that nsP3 is able to interfere with ARTD10 function in infected cells. We propose therefore that the nsP3 macro domain directly interacts with one or more ADP-ribosylated proteins involved in the NF-κB pathway, likely candidates being NEMO and/or ARTD10. However, given the previous observation that miR-146a is also involved in the regulation of the NF-κB pathway by CHIKV, we believe it is highly likely that CHIKV, like many other viruses, possesses multiple mechanisms to control this key signalling pathway.

Analysis of the nsP3 macro domain mutant panel in the context of infectious CHIKV revealed that there was little correlation between the virus growth phenotype and the effect on the NF-κB pathway (Fig 7). Thus we conclude that, at least in the range of cells tested, inhibition of NF-κB activation is not required for efficient production of infectious virus. However, it was interesting to note that most mutants exhibited cell type specific phenotypes, for example production of virus in Huh7 cells seemed particularly dependent on macro domain function as 4 mutants failed to grow in these cells, of these 3 (D10A, G32A and V113A) were able to grow in other cell types. The growth of some mutants was not due to reversion to wildtype as, at least for G32A, T111A and Y113A we did not observe any reversion to wildtype in C2C12 or U4.4 cells (data not shown). However, due to low titres we were unable to RT-PCR and sequence the other mutants.

In conclusion, we show here that one of the roles of the CHIKV nsP3 macro domain is to contribute to the regulation of a critical antiviral factor, the transcription factor NF-κB. This observation leads to a number of key questions for the future: for example is this function conserved in the other viral macro domains? What are the other targets for the ADP-ribosylhydrolase activity of these domains? And indeed, why is the macro domain only present in a small subset of positive-strand RNA viruses? What is clear is that we have only just begun to understand the role of this domain in the lifecycle of these viruses.

## Author Statement

### Author contributions

GCR performed the experiments, MH supervised the study with assistance from NJS. All authors were involved in writing the manuscript.

### Conflicts of Interest

The authors declare that there are no conflicts of interest.

### Funding Information

GCR was a PhD student funded by the Biotechnology and Biological Sciences Research Council (BBSRC) White Rose PhD programme in Mechanistic Biology (grant number BB/F01614X/1). This work was supported by a Wellcome Trust Investigator Award to MH (grant number 096670).

## Acknowledgements

We thank Andres Merits (University of Tartu, Estonia) for CHIKV replicon and infectious virus constructs, anti-nsP3 antibody and helpful advice. We also thank Geoffrey Smith (University of Cambridge) for the vaccinia B14 expression construct, and Andrew Macdonald (University of Leeds) for the NF-κB responsive reporter construct. The authors acknowledge Carsten Zothner for the technical assistance with BSL3 studies.

---

**Supplementary Fig 1.**
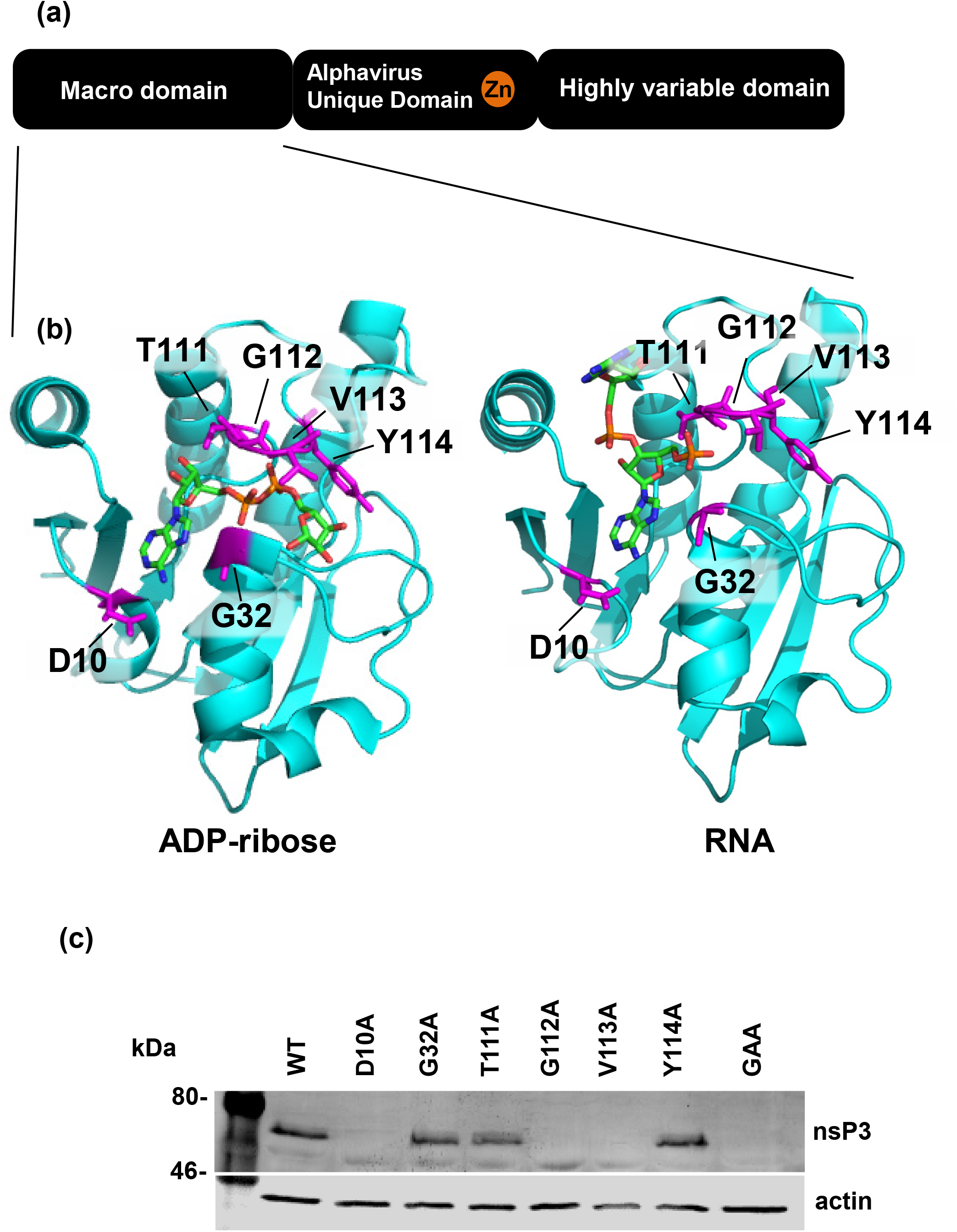
**(a)**. Schematic of the 3 domain structure of nsP3. **(b)**. Structure of the CHIKV macro domain in complex with ADP-ribose or RNA, showing position of mutated residues. (PDB accession numbers 3GPO and 3GPQ respectively^15^). **(c)**. Western blot showing expression of nsP3 by CHIKV macro domain mutant viruses in BHK-21 cells.

**Supplementary Fig S2.**
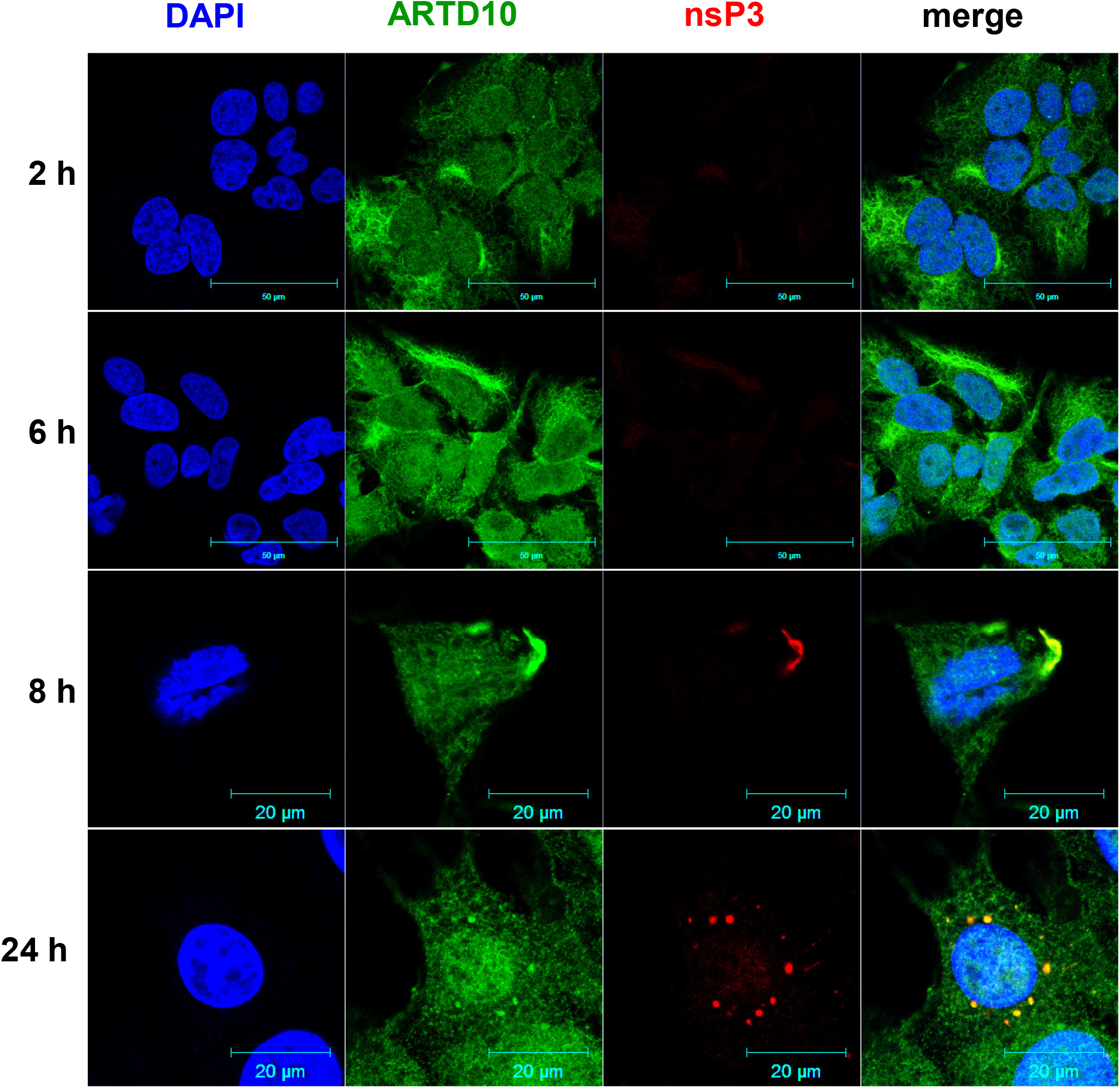
Huh7 cells infected with CHIKV were fixed and stained with antibodies to ARTD10 and nsP3 at the indicated times after infection.

